# Controlling for polygenic genetic confounding in epidemiologic association studies

**DOI:** 10.1101/2024.02.12.579913

**Authors:** Zijie Zhao, Xiaoyu Yang, Jiacheng Miao, Stephen Dorn, Silvia H. Barcellos, Jason M. Fletcher, Qiongshi Lu

**Author notes:** To whom correspondence should be addressed: Dr. Qiongshi Lu.

## Abstract

Epidemiologic associations estimated from observational data are often confounded by genetics due to pervasive pleiotropy among complex traits. Many studies either neglect genetic confounding altogether or rely on adjusting for polygenic scores (PGS) in regression analysis. In this study, we unveil that the commonly employed PGS approach is inadequate for removing genetic confounding due to measurement error and model misspecification. To tackle this challenge, we introduce PENGUIN, a principled framework for polygenic genetic confounding control based on variance component estimation. In addition, we present extensions of this approach that can estimate genetically-unconfounded associations using GWAS summary statistics alone as input and between multiple generations of study samples. Through simulations, we demonstrate superior statistical properties of PENGUIN compared to the existing approaches. Applying our method to multiple population cohorts, we reveal and remove substantial genetic confounding in the associations of educational attainment with various complex traits and between parental and offspring education. Our results show that PENGUIN is an effective solution for genetic confounding control in observational data analysis with broad applications in future epidemiologic association studies.

## Introduction

Identifying environmental risk factors for health outcomes can have important implications on public health and precision medicine^1-3^. However, epidemiologic associations are often confounded by genetics in observational studies^4,5^. For example, the association between maternal smoking during pregnancy and child psychopathology can be largely explained by the shared genetic variations of mother and child^6^. Genome-wide association studies (GWAS) in the past two decades have identified tens of thousands of single-nucleotide polymorphisms (SNPs) for human complex traits^7-9^. We now understand that most human traits are highly polygenic and genetic associations are often shared across many phenotypes^10-13^. Failing to adjust for such pervasive genetic confounding can lead to bias in effect size estimation and misinterpretation of association results. Consequently, medical interventions and clinical decisions based on observational associations without genetic confounding control may fail. For example, recent studies have shown that the associations previously found between vitamin D level and both type-I diabetes and multiple sclerosis are confounded by genetic variants altering both vitamin D metabolism and disease risk^14,15^. Therefore, vitamin D supplementation and related medical interventions may not effectively prevent the onset or progression of these health issues.

However, it is challenging to control for genetic confounding in observational studies. Although the confounding effect of any individual genetic variant is usually weak, the total number of confounding variants can be large^7,16^. These individually minimal but cumulatively non-negligible confounding effects pose a major challenge to study design and analytical approaches^17-19^. Classic approaches use monozygotic twins^20-22^, and more recently, dizygotic twins and full siblings^23-25^, who share a large fraction of genotypes (and environment) to control genetic (and environmental) confounding. While these studies are generally considered more robust compared to analysis based on unrelated individuals in population cohorts, they often have limited sample sizes and insufficient statistical power^20,26^. In comparison, studies leveraging large biobank cohorts of mostly independent samples have superior statistical power^27-30^. These studies either do not adjust for genetic confounding at all^31-33^, or use molecular genetic data to adjust for confounding^34,35^. Some studies control individual SNPs as covariates^36,37^. But due to the polygenic nature of genetic confounding, recent studies often first aggregate many SNP effects into polygenic scores (PGS), then adjust for these scores in association analysis^29,38,39^. These studies have several major issues. First, empirical PGS calculated from GWAS data are very noisy estimates of true genetic risks^17,18^. As we demonstrate below, adding these noisy scores as covariates does very little to control for genetic confounding. Second, using PGS as covariates implies strong assumptions on the confounding mechanism; that is, researchers must select which PGS controls to use (and therefore which ones not to use). It is inherently a hypothesis-driven approach assuming that a particular genetic trait drives all the confounding. It is unclear in the current literature whether PGS of the exposure, the outcome, or some other traits should be adjusted for in the analysis^27-29,38^. Third, PGS-based methods require access to an individual-level cohort with exposure, outcome, and genetic variables. In addition, this study cohort needs to be independent from GWAS samples in which PGS models are trained^18,40^. In practice, separate GWAS and individual-level genotype and phenotype datasets can be difficult to acquire, thus limiting the utility of these methods.

Here, we introduce PENGUIN (PolygENic Genetic confoUnding INference), a variance component analytical framework to control for genome-wide confounding effects in epidemiologic association studies. Using the PENGUIN framework, we showcase a strategy to fully control for polygenic genetic confounding without hypothesizing any specific confounder traits. We also demonstrate that PENGUIN does not require GWAS samples and association study samples to be independent. In fact, we show that under some general conditions, there is no need to use individual-level data at all – having GWAS summary statistics alone is sufficient to produce the association results of interest while adjusting for genetic confounding. Through extensive simulation analyses, we demonstrate that PENGUIN provides well-calibrated confidence intervals and type-I error rates under all settings. We apply our approach to estimate the contribution of educational attainment (EA) on a variety of complex phenotypes in the UK Biobank^41^ (UKB) and evaluate the effect of maternal/paternal EA on offspring EA in the Health and Retirement Study^42^ (HRS).

## Results

### Method overview

PENGUIN is a principled approach to controlling for polygenic genetic confounding with minimal requirements on the input data and confounding mechanism. The PENGUIN framework contains three steps (**Figure 1A**). First, it estimates the heritability of exposure and the genetic covariance between exposure and outcome using GWAS summary statistics^11,12^. Second, PENGUIN calculates the phenotypic covariance between exposure and outcome using either individual-level phenotype data or GWAS summary statistics. Finally, it estimates the unconfounded exposure effect on the outcome. We refer to the version of PENGUIN that only relies on GWAS summary statistics as PENGUIN-S. Details of the statistical framework are described in **Methods**. For comparison, we also illustrate a common approach that uses PGS to adjust for genetic confounding (**Figure 1B**). This approach requires an individual-level dataset as well as a separate GWAS without any sample overlap, which is a key data constraint PENGUIN is able to remove (**Methods**).

**Figure 1.**
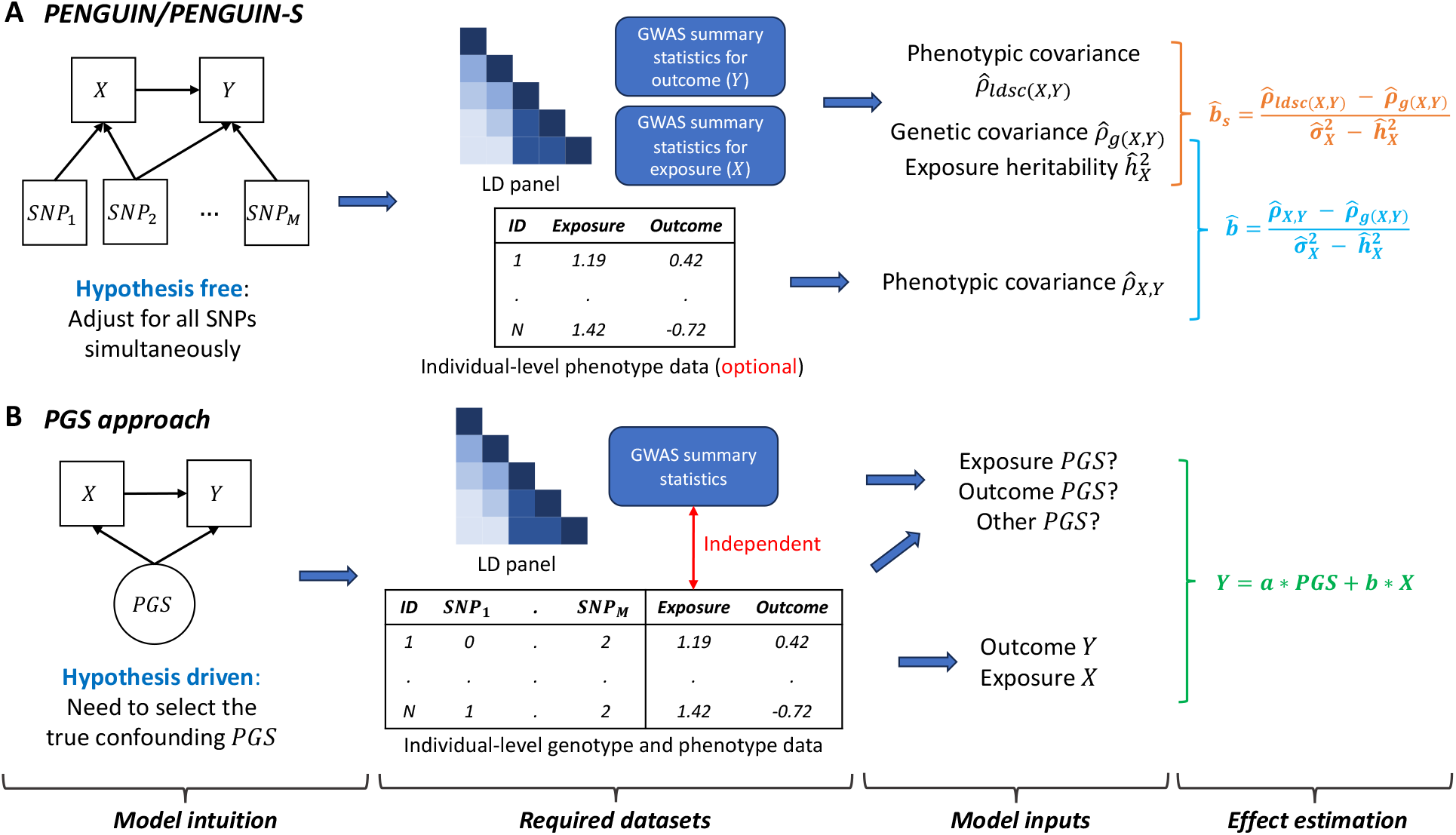
Workflow for removing genetic confounding effects in observational data analysis. (**A**) Both PENGUIN and PENGUIN-S quantify the exposure-outcome association while adjusting for all SNPs in the analysis. PENGUIN uses both summary-level GWAS data and individual-level phenotype data as inputs. PENGUIN-S employs a summary-statistics-based approach without relying on any individual-level data. (**B**) Methods using PGS as input hypothesize that the genetic confounding can be quantified by PGS of a particular trait. These methods demand independent GWAS summary statistics and individual-level genotype and phenotype data for PGS calculation and association analysis

### Simulation results

We conducted extensive simulations using genotype data from the Wellcome Trust Case Control Consortium^43^ (WTCCC) to validate the performance of PENGUIN. After quality control, we included 15,567 samples and 292,470 variants in the simulation study. We simulated the outcome (i.e., Y) and exposure variables (i.e., X) under diverse settings by varying the sparsity of causal genetic effects, magnitude of genetic confounding effects, and the true association effect between exposure and outcome (**Methods**). We compared PENGUIN and PENGUIN-S with the common approach that adds PGS of exposure/outcome as a covariate in regression (denoted as PGSX/PGSY hereafter). We also compared with an approach that applies structural equation modeling to correct for measurement error in the PGS covariate^18^. We refer to this as GsensX/GsensY based on whether the exposure or outcome PGS was adjusted for. For fair comparison, we divided the study sample into a training set (N=12,000) for GWAS, a testing set (N=3,000) for exposure-outcome association estimation, and an additional reference set (N=567) for linkage disequilibrium (LD) estimation. For PGSX/PGSY and GsensX/GsensY approaches, we computed clumping and thresholding (C+T) PGS on the testing set. For PENGUIN, we fitted LD score regression^11,12^ using LD scores constructed from the reference set. In total, we investigated 13 different simulation settings and repeated each setting 100 times. Details on the simulation design are described in **Methods** and **Supplementary Table 1**.

Our method provided unbiased estimates of exposure-outcome associations with well-controlled type-I error and confidence interval coverage in all settings (**Figure 2A-B** and **Supplementary Figures 1-6**; **Supplementary Tables 2-4**). In addition, results from PENGUIN-S without using any individual-level testing data were highly consistent with PENGUIN, showcasing the performance and validity of this full-summary-statistics approach. Meanwhile, marginal regression between the outcome and exposure resulted in highly biased coefficient estimates when genetic confounding was present. Adjusting for empirical PGS in regression (i.e., PGSX and PGSY) barely removed any genetic confounding effects and led to highly biased association results. In comparison, GsensX corrected for measurement error in PGS and provided unbiased association estimates; however, it had poorly calibrated confidence intervals which led to a 3.8-fold inflation of type-I error. We note that GsensY showed substantial biases compared to true association values, highlighting the importance of properly choosing the genetic score in PGS-based approaches. We provide more discussions on this issue in later sections.

**Figure 2.**
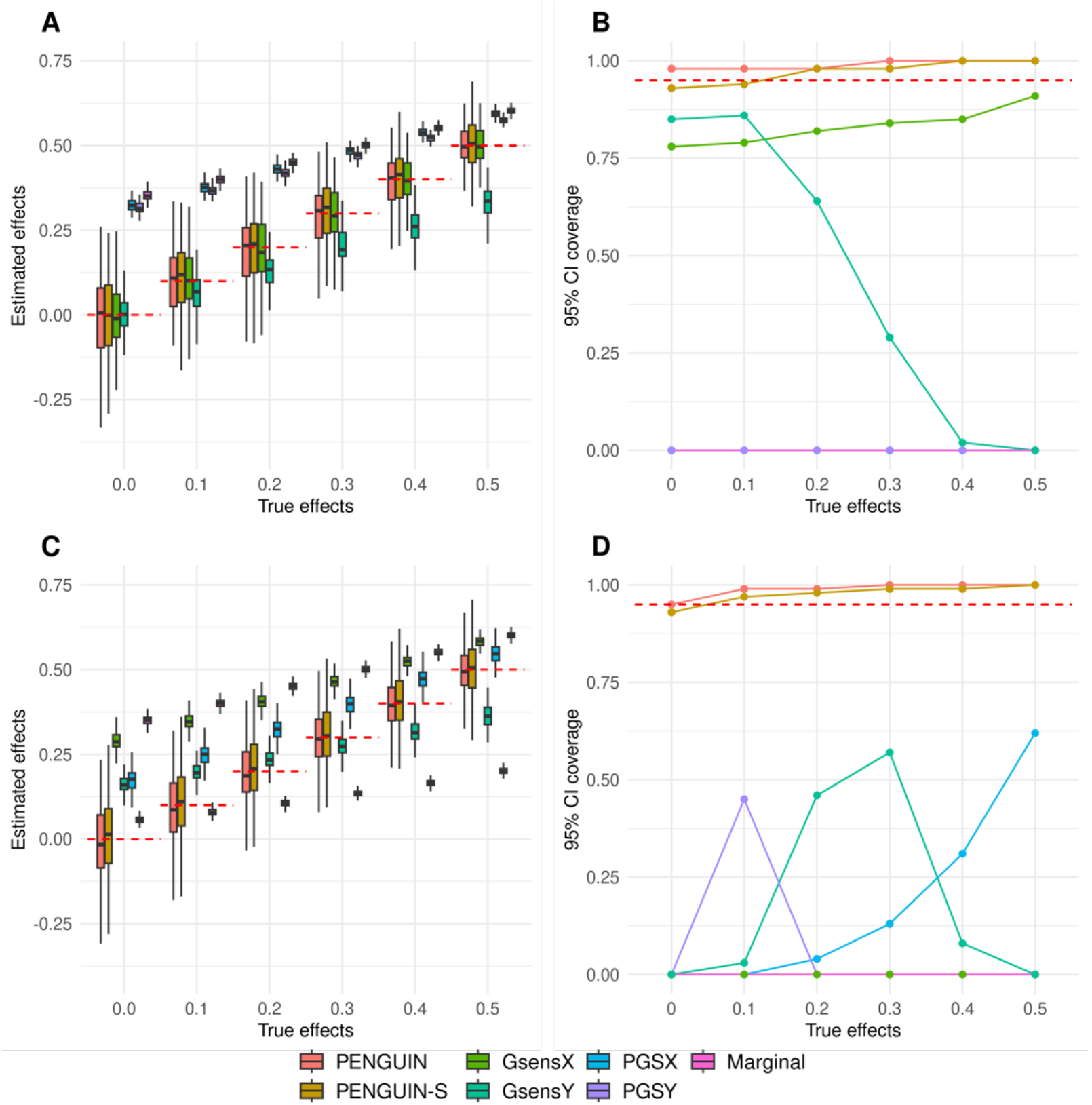
Simulation results. (**A** and **C**) Exposure-outcome associations estimated by different methods across 100 replications. (**B** and **D**) 95% confidence interval coverage for each method across 100 replications. GWAS summary-level data and individual-level testing dataset are independent in **A**-**B** (0% sample overlap) while all testing samples are included in the GWAS data in **C**-**D** (100% sample overlap). Y-axis: exposure effects for **A** and **C** and 95% coverage for **B** and **D**; X-axis: true exposure effect size. Red dashed lines are true effects in **A** and **C** and 95% coverage threshold in **B** and **D**. Across settings shown in this figure, proportion of causal variants is 0.1% and true genetic confounding effects are the entire genetic component for the exposure. Detailed simulation settings are described in **Supplementary Table 1**. Results for other simulation settings are summarized in **Supplementary Tables 2-4** and **Supplementary Figures 1-6**.

Next, we investigated the robustness of all methods under the presence of sample overlap between GWAS and the exposure-outcome study cohort. For each previous simulation setting, we considered two more scenarios where 50% or 100% of samples in the testing data are also included in the training GWAS dataset. In these scenarios, only PENGUIN and PENGUIN-S provided robust and accurate estimation of true exposure effects (**Figure 2C-D**). As expected, methods that require PGS as inputs became impossible to use when independent individual-level samples are unavailable, showing substantial estimation biases and poor confidence interval coverages. Simulation results for the remaining settings are summarized in **Supplementary Figures 1-6** and **Supplementary Tables 2-4**.

### Associations of education with 26 complex traits in UKB

Education is known to be associated with many later life health outcomes^21,23,44,45^. However, studies have also reported substantial pleiotropy of genetic variants associated with EA and various complex traits^46-48^. We applied PENGUIN to systematically evaluate the effect of EA on a variety of health and social outcomes in UKB^41^ (**Methods**). Following recent studies^48,49^, we included 26 phenotypes including 15 quantitative traits and 11 binary traits in this analysis. The total sample sizes of these phenotypes range from 2,409 to 406,803. Information on each phenotype, corresponding individual-level and summary-level datasets, and their heritability and genetic correlations/covariances with EA are summarized in **Supplementary Tables 5-6** and **Supplementary Figure 7**. We also included results from marginal regression and PGSX to show the importance and effectiveness of genetic confounding control.

We found substantial reductions in EA effects on many traits after controlling for genetic confounding using PENGUIN (**Figure 3A-B**; **Supplementary Table 7**). In comparison, the PGS approach had minimal impact on EA effect estimation. Using marginal regression estimates as the baseline, PENGUIN and PGSX achieved median reductions of 19.3% and 3.3% in EA effects on all traits (**Supplementary Figure 8**). The most substantial EA effect reductions were observed for bipolar disorder and ever smoking. We also compared PENGUIN with PENGUIN-S and obtained nearly identical EA effect estimates (cor=0.996) (**Figure 3C**; **Supplementary Figure 9**; **Supplementary Table 8**).

**Figure 3.**
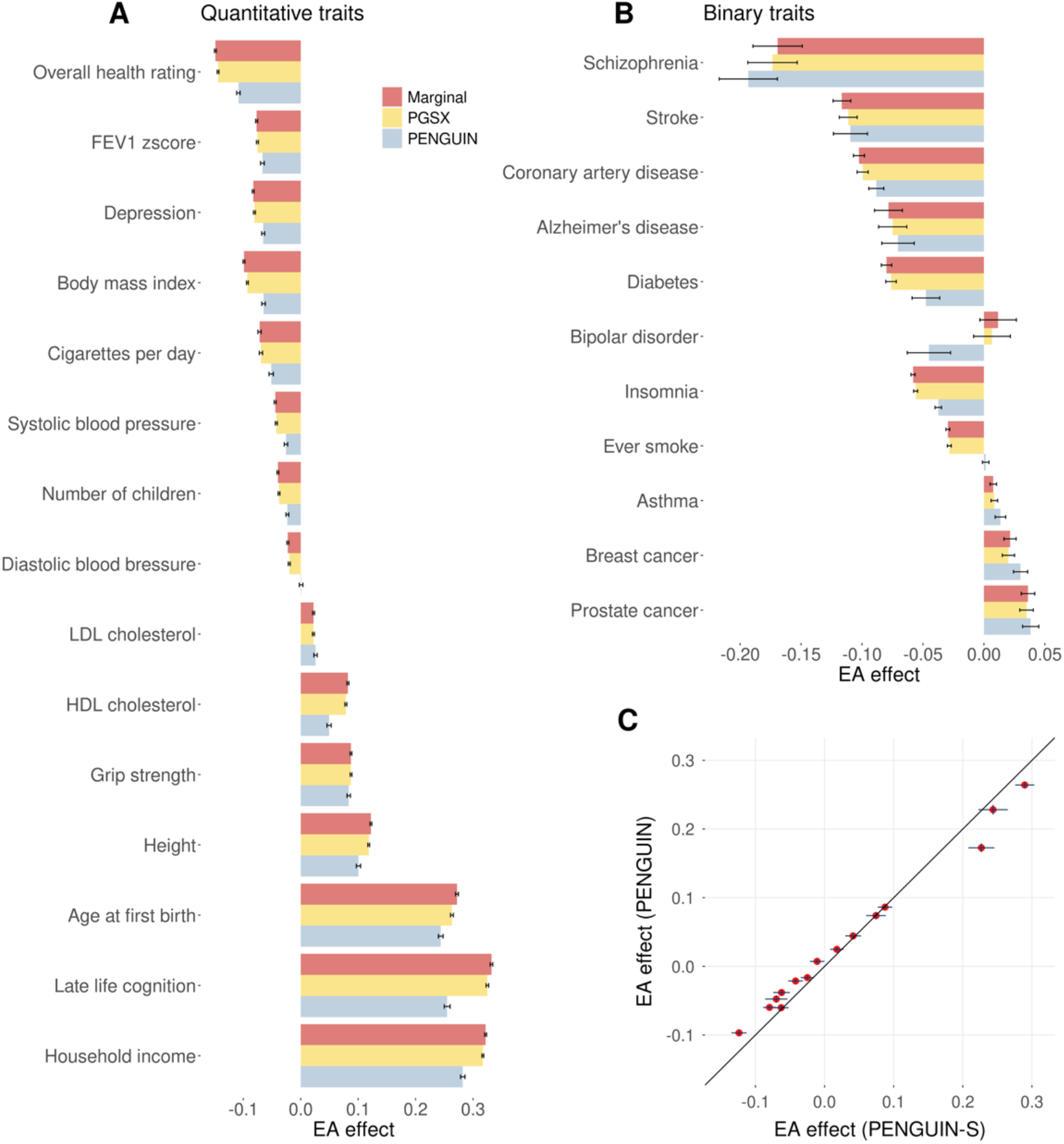
EA effects on various complex traits in UKB. (**A**-**B**) Comparing EA effects on complex traits estimated by PENGUIN, PGSX, and marginal regression. Effect size point estimates and standard errors for quantitative and binary traits are shown in panels **A** and **B**, respectively. Y-axis: outcome trait; X-axis: EA effect size estimates (per SD change in EA). (**C**) Comparison between EA effects estimated by PENGUIN and PENGUIN-S for 15 quantitative traits. Y-axis: PENGUIN estimates; X-axis: PENGUIN-S estimates. The black solid line is the diagonal line. Standard error bars from both approaches are shown for all data points. Full association results are summarized in **Supplementary Tables 7-8** and **Supplementary Figures 8-9**.

Education is known to associate with higher later life cognition^23,50^. However, analyses focusing on this association using observational data did not always adjust for genetic confounding^51,52^. PENGUIN estimated a positive effect of EA on cognition in UKB: one SD increase in EA is associated with a 0.25-SD increase in cognition, which reduces the association effect by 23% and 21.5% compared to marginal regression and PGSX, respectively. This finding is consistent with the attenuated positive effect of EA on cognition observed in sibling studies and causal studies^23,49^. It also highlights again that a simple PGS approach is insufficient to account for genetic confounding effects despite the large GWAS sample size for EA.

Several traits showed amplified associations with EA after controlling for genetic confounding. For example, PENGUIN increased the protective association of EA on schizophrenia (SCZ) risk by 14.2%. Similarly, we found a null but positive EA association with bipolar disorder when genetics was not controlled for, but PENGUIN identified a significant protective EA effect on bipolar disorder (BD) (**Figure 3B**). Effect size amplification is possible when genetic confounding and EA effect operate in the opposite directions (**Methods**), which can be seen in the case of SCZ (i.e., EA-SCZ genetic covariance=0.002, phenotypic covariance=-0.169). Or, in the case of EA-BD association, genetic covariance had the same sign but a higher magnitude compared to phenotypic covariance (EA-BD genetic covariance=0.054, phenotypic covariance=0.012). Our positive genetic correlation estimates for EA with SCZ and BD are consistent with what has been reported in the literature, but interestingly, intelligence is known to have negative genetic correlations between with both SCZ and BD^53^. Therefore, we hypothesize that the genetic confounding of EA-SCZ and EA-BD associations may be driven more by the non-cognitive component of EA. To assess this hypothesis, we partitioned EA genetics into orthogonal cognitive and non-cognitive components using the GWAS-by-subtraction approach^54^ to study their each respective confounding effect. We found positive genetic correlations between the non-cognitive component of EA and both SCZ and BD (**Figure 4A-B**; **Supplementary Table 9**). When controlling for the non-cognitive component of EA genetics, we found significant negative associations of EA on SCZ and BD (**Figure 4C-D**). In comparison, the cognitive component of EA genetics contributed much less to the overall genetic confounding. Together, two independent EA genetic components demonstrated opposite directions and different degrees of confounding effects on the association between EA and two psychiatric disorders (**Supplementary Figure 10**). In addition to SCZ and BD, we also found amplified EA effects on higher risks of breast cancer and prostate cancer which may be explained by greater exposure to risk factors^55,56^, higher screening rate and lower mortality rate^57^ among better educated individuals. Therefore, the seemingly damaging effect of EA on these health outcomes should be interpreted with caution, especially when non-genetic confounders have not been fully accounted for.

**Figure 4.**
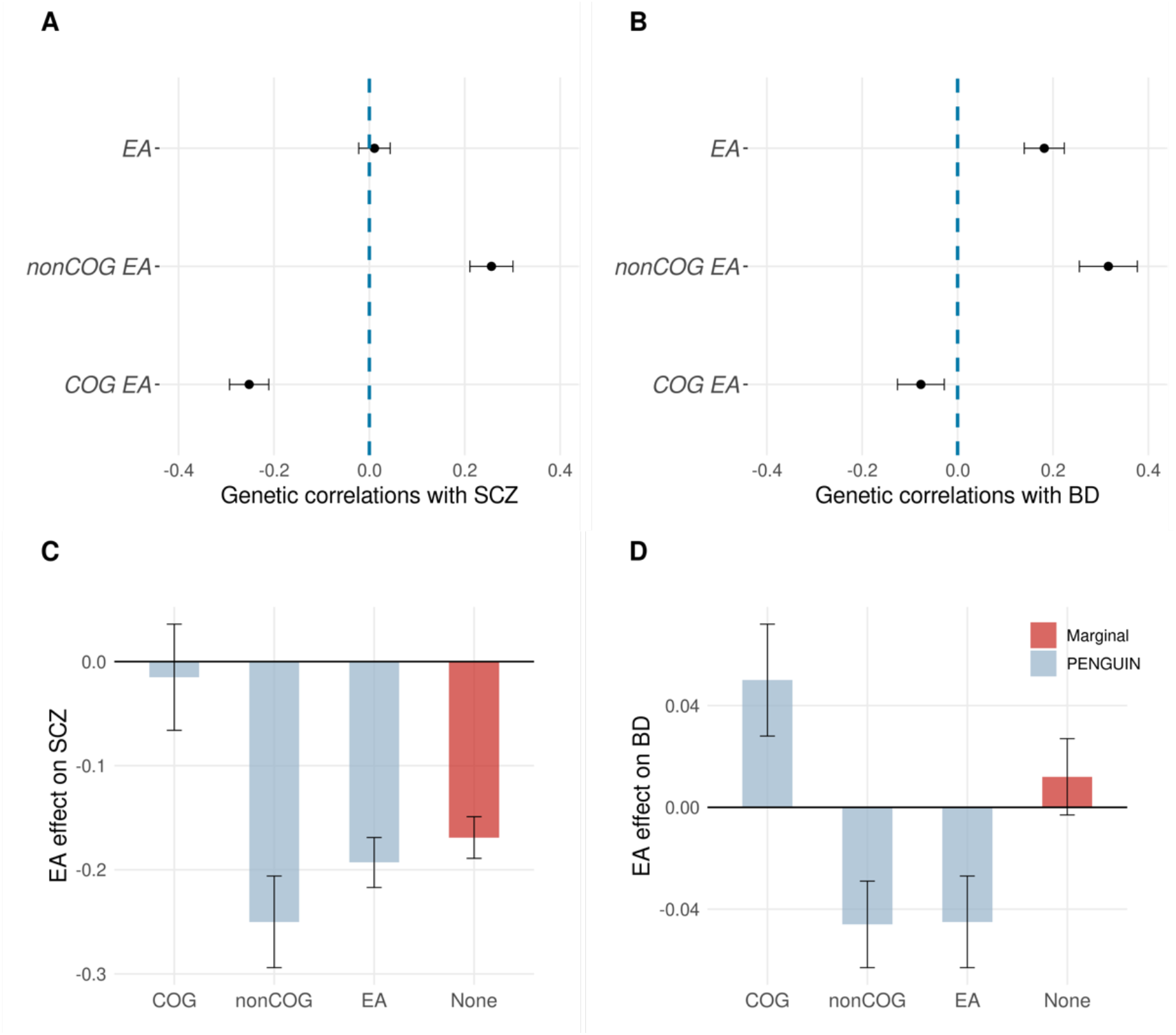
Associations of EA with SCZ and BD. (**A**-**B**) Genetic correlation between SCZ/BD and cognitive/non-cognitive/full EA genetic components. Y-axis: EA genetic components; X-axis: genetic correlation. (**C**-**D**) Estimated EA effects on SCZ and BD after controlling for cog/non-cog/full EA genetic components. Y-axis: effect size estimates (per SD change in EA); X-axis: adjusted EA genetic components. Standard error bars are shown for all data points. Full association results, heritability and genetic covariance/correlation estimates are summarized in **Supplementary Table 9**. SCZ: schizophrenia; BD: bipolar disorder.

### Estimating parental effects on offspring EA

Understanding how parental behavior influences children’s developmental outcomes is of broad interest in several research fields^58-60^. However, observed associations between parental and offspring outcomes can be confounded by genetics due to Mendelian genetic transmission^61-63^. Here, we expanded the PENGUIN framework to control for confounding due to inter-generational genetic transmission (**Methods**), and estimated the association between parental EA and offspring EA in HRS^42^ (N=12,068; **Methods**).

We obtained significant positive effects of both maternal and paternal EA on offspring EA, but these effects were weakened after proper control of genetic confounding (**Figure 5** and **Supplementary Table 10**). Maternal and paternal EA effects reduced by 14.7% and 14.4%, respectively, after removing genetic confounding. Simply adding EA PGS in the regression only resulted in reductions of 5.8% and 6.1% in effect estimates. As a secondary analysis, we allowed genetic associations with paternal and maternal EA to be different (**Methods**), and observed even more substantial reductions in maternal (22.2%) and paternal (19.1%) effects on offspring EA (**Supplementary Figure 11**).

**Figure 5.**
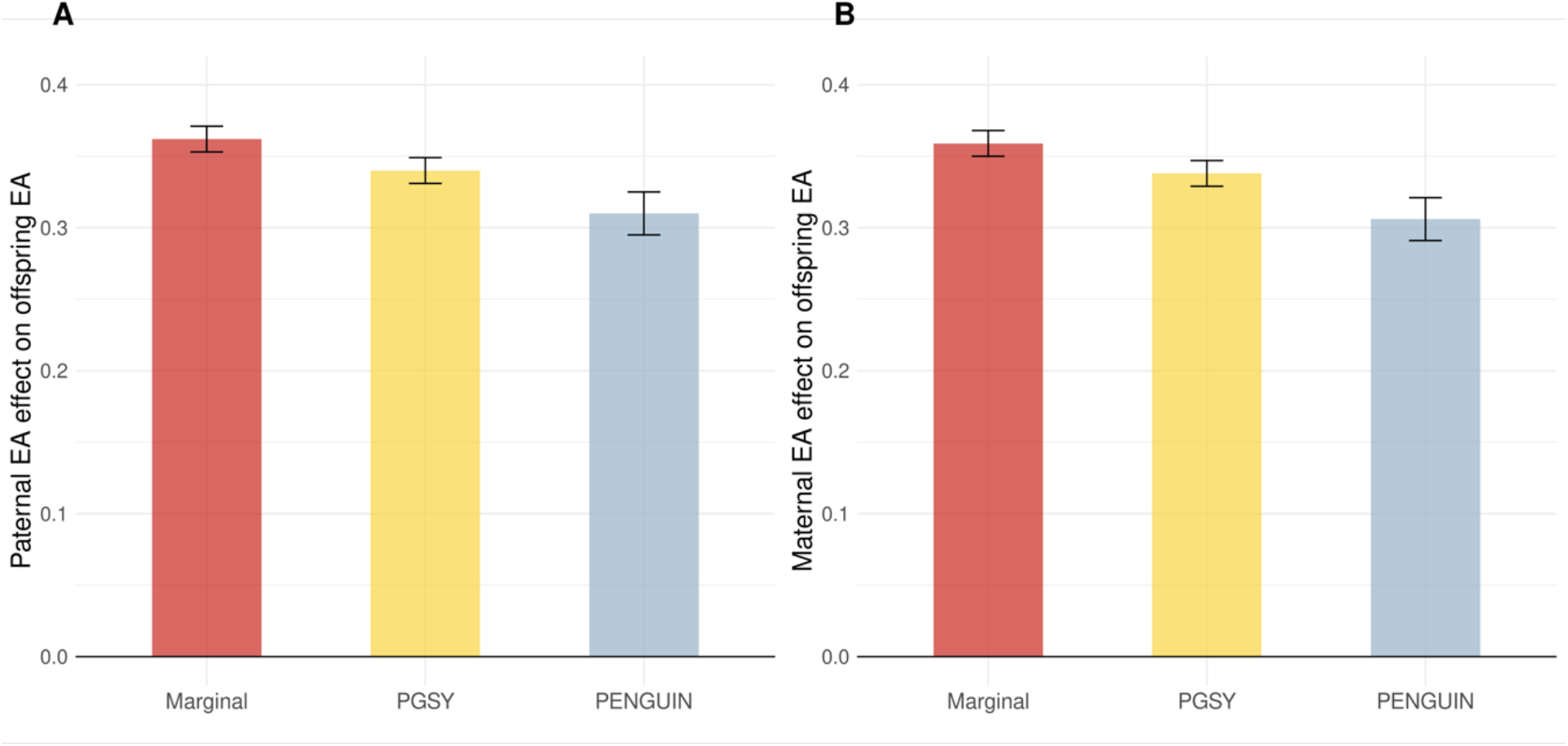
Effects of parental EA on offspring EA. Paternal EA (**A**) and maternal EA (**B**) effect sizes are estimated from PENGUIN, PGSY, and marginal regression. Y-axis: EA effect size estimates (per SD change in parental EA); X-axis: association approaches. Additional association results are summarized in **Supplementary Table 10** and **Supplementary Figure 11**.

## Discussion

Accurate estimation of unconfounded association between exposure and outcome variables is a foundational task in epidemiology and social sciences. Due to widespread sharing of genetic associations across complex human traits, it is crucial to fully control for genetic confounding that may otherwise lead to incorrect conclusions in observational data analysis^12,13^. To address this challenge, we introduced PENGUIN, a principled framework to effectively control for genetic confounding using GWAS summary statistics. Through extensive simulations, we demonstrated that both PENGUIN and PENGUIN-S provide accurate effect size estimation with well-calibrated confidence intervals. Applying PENGUIN to multiple large cohorts, we showed that controlling for genetic confounding will sometimes substantially shift the association effect estimates between exposures and outcomes.

Our study brings multiple advances to the field. First, most existing studies implicitly or explicitly rely on assumptions on the mechanisms of genetic confounding and need to specify the traits whose PGS are included in the regression model^27-29^. The validity and interpretation of these association estimates depend heavily on correctly specifying the true confounding PGS in the model. But in practice, these assumptions can be easily violated. In comparison, PENGUIN bases its inference framework on a high-dimensional regression model which accounts for all genetic variants’ confounding effects without implying any structure on these effects across SNPs. Second, PENGUIN is currently the only method that provides valid, genetically-unconfounded inference based on population cohorts. We have demonstrated that the commonly used PGS approach is ineffective for genetic confounding control due to noises in empirical PGS even under biobank-scale GWAS sample sizes. Measurement error correction can somewhat reduce this issue but leads to inflated type-I errors and poorly calibrated statistical inference. While the cause for the statistical mis-calibration still remains to be investigated, we provide an alternative solution using a variance component analytical framework. PENGUIN uses a different framework based on variance and covariance components (i.e., for heritability and genetic covariance estimation) and is therefore immune from any measurement error in PGS^11,12^. We have previously explored related ideas in the context of quantifying gene-environment interaction^64^. This work can be viewed as an extension of our previous work into the new application of removing polygenic genetic confounding. Third, PENGUIN is highly flexible with its input data. The classic PGS approach and its measurement-error-corrected extension require GWAS summary statistics independent from the association study samples to avoid overfitting. This can create challenges in practice, since publicly available GWAS statistics datasets often include samples from many smaller cohorts through meta-analysis. PENGUIN is the only method that can be applied to datasets with overlapping samples without biasing association findings. Finally, PENGUIN-S further reduces the data requirement without compromising the validity of association results. When the GWAS summary-level datasets for exposure and outcome share a substantial number of individuals, we have demonstrated that individual-level genotype and phenotype data will no longer be needed for the analysis. With the increasing availability of GWAS summary statistics in the public domain, we expect PENGUIN-S to become an important tool with broad applications in future studies.

Applying PENGUIN to UKB, we found evidence for pervasive genetic confounding in the associations between EA and health and social outcomes. After controlling for confounding, we found substantial attenuation of EA effect on other traits. For example, genetics explains about 20% of the association between EA and later life cognition. SCZ and BD are two psychiatric disorders marked by impaired cognitive function^65,66^. Interestingly, BD has been shown to positively correlate with higher EA on both the phenotypic and genetic levels^67-69^. We confirmed these correlations based on UKB data, and more importantly, we found that positive correlation between EA and BD was driven by non-cognitive component of EA, which was the genetic confounder that masked the protective effect of EA against BD. We also found similar results for the EA-SCZ association. Finally, vast literature suggests that parental EA can positively affect offspring EA by stimulating development of both cognitive and non-cognitive skills through genetic inheritance or family environment^59,63,70,71^. In a study that tried to disentangle genetics from environment, Conley et al. concluded that genetics accounted for one sixth of the correlation between offspring and parental EA, but no significant difference was found in parental EA effects calculated with and without adjusting for EA PGS^63^. Using PENGUIN, we found a much larger attenuation of parent-offspring EA association after controlling for genetic confounding. These results highlight the superior performance of PENGUIN compared to the classic PGS approach in adjusting for genetic confounding, and showcase the importance of accounting for inter-generational genetic transmission in future studies of parental effects on child cognitive outcomes.

Our study has some limitations. First, we did not attempt to adjust for unmeasured non-genetic confounding in our analyses. For this reason, we compared PENGUIN with the PGS approach but not twin or sibling-based analysis since those study designs also control for environmental confounding. Partly due to this, we do not make any causal claims. The choice of analytical approach in future studies should depend on the study aims. If the main goal is to remove genetic confounding in observational data analysis using independent individuals, PENGUIN is an ideal strategy. We recommend researchers to thoroughly check for non-genetic confounding in addition to PENGUIN analysis and always interpret association results with caution. Second, PENGUIN substantially reduced the estimates of EA effects on many health and social outcomes in our analysis, but it remains challenging to validate these estimates in real data due the lack of ground truth. Third, because PENGUIN bases its framework on a high-dimensional linear regression model of additive genetic effects, non-linear genetic confounding and gene-gene or gene-environment interaction^64^ confounding, if exist, can still bias association results. For binary outcomes, it is also of interest to extend PENGUIN to generalized linear model or calculate odds ratios of exposures in the future^72^. Further, we only included individuals of European descent in the current analyses. While our approach is applicable to datasets of any genetic ancestry, heritability and genetic covariance estimators can be sensitive to the choice of LD reference data and computational tools^73-75^, which could alter downstream association results and interpretation. Lastly, the current PENGUIN framework is only applicable for investigating the association between a pair of exposure and outcome variables while accounting for their shared genetic basis. It remains a future direction to generalize PENGUIN so it can estimate the effects of multiple exposures on the outcome of interest while controlling for genetic confounding.

In summary, in this work we introduced an effective and efficient framework for genetic confounding control in observational association studies. As a variance component-based framework, our approach is immune from assumptions and measurement errors of PGS and is robust to sample overlap between multiple input datasets. As a full-summary-statistics technique, PENGUIN-S further reduces data requirement and produces highly consistent results. Together, PENGUIN and PENGUIN-S are a highly flexible framework that can have broad applications in future observational data analysis.

## Methods

### PENGUIN framework

PENGUIN is based on the following linear model for exposure and outcome variables *X* and *Y*. For simplicity, we assume that both *X* and *Y* are standardized with mean 0 and variance 1:

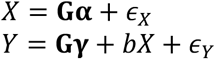

where **G** = [*G*_1_, *G*_2_, …, *G*_M_] is a random vector of the genotype of all SNPs and *M* is the total number of SNPs; **α** = [α_1_, …, α_M_]^T^ and **γ** = [*γ*_1_, …, *γ*_M_]^T^ are random SNP effects with mean 0 for *X* and *Y*, respectively; *∈*_X_ and *∈*_Y_ are centered non-genetic effect terms with positive and finite variances. We assume *∈*_X_, *∈*_Y_, and **G** to be mutually independent. The main parameter of interest is b, which quantifies the association of exposure *X*with outcome *Y*. No additional assumption on the distributions of **α**, **γ**, *∈*_X_, and *∈*_Y_ was made, and **α** and **γ** can be either independent or correlated.

While it seems straightforward to estimate *b* through a regression of *Y* on *X* with **G** added as covariates, this could be impossible in practice due to the large dimensionality of SNP data (*M* could be several million). However, we note that fitting a regression model *Y*∼ *X*+ **G** is equivalent to first regressing out **G** from both *X* and *Y* and then fitting a regression between residualized *Y* and *X*. Therefore, with unlimited sample size, the exposure effect can be obtained by:

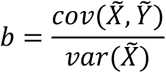

Where 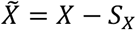 and 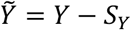, and ***S***_X_ and ***S***_Y_ are the additive genetic components underlying *X* and *Y*, respectively (i.e., ***S***_X_ = **Gα** and ***S***_Y_ = **G(**b**α** + **γ)** according to the model above). Based on this, we have:

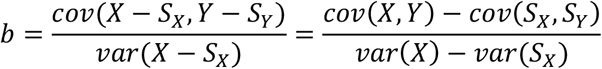

It is straightforward to estimate *cov*(*X, Y*) and *var*(*X*) from phenotypic data. Further, *cov*(***S***_X_, ***S***_Y_) and *var*(***S***_X_) are the genetic covariance between *X* and *Y* and heritability of *X*, respectively. Thus, in practice, estimate the exposure effect *b* by:

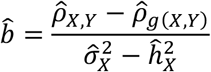

Where 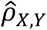 and 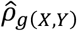 are the estimated phenotypic and genetic covariance between *X* and *Y*, and 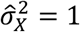 and 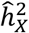 are the sample variance and estimated heritability of *X*. 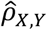 is estimated from individual-level data of *X* and *Y*; 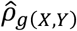 and 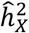 can be estimated using GWAS summary statistics. We used LD score regression to obtain both 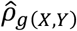 and 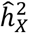 throughout this study^11,12^.

### Interval estimation and hypothesis testing

By central limit theorem, both sample covariance 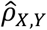 and sample variance 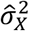 converge in distribution to normal random variables, given the typically large GWAS sample size:

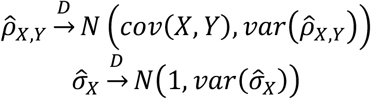

Empirically, we employ the following approximations for 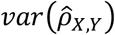 and 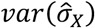 based on normal theory^76^ when analyzing complex traits:

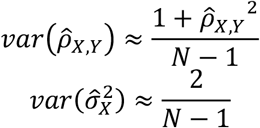

Let 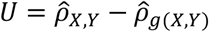 and 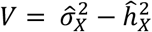 denote the numerator and denominator in 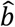, respectively. Then, we have:

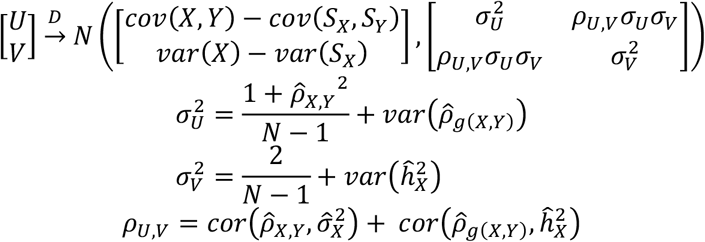

where *N* is the individual-level sample size of exposure *X* and outcome *Y*, and 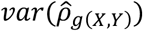 and 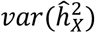 are sampling variance of genetic covariance and heritability estimators from LD score regression based on block Jackknife^11,12^. In practice, we ignore the covariance term *ρU*,V^*σ*^*U*,^*σ*^Vsince *ρU,V* is close to 0 empirically^77^. Then, by delta method, 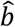 converges in distribution to a normal variable with variance:

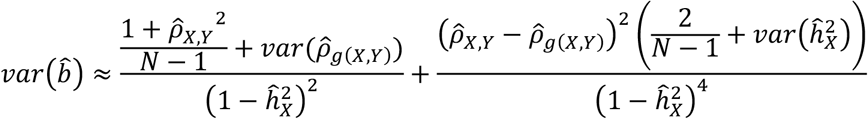

We can use Z statistics and the corresponding p-value to evaluate the null hypothesis *H*_*0*_: *b* = *0*.

Note that we assume pairwise independence between LD score regression estimators (i.e., 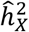 and 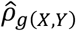) and estimators based on individual-level phenotype data (i.e., 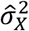 and 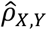) when deriving the distribution of 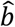. This is true when GWAS samples are independent from the dataset for *X* and *Y*. When two datasets share a subset of individuals, this assumption is violated. Nonetheless, we demonstrate in simulations that violating this assumption has minimal influence on the performance of PENGUIN even when there is substantial sample overlap between two datasets (**Figure 2**; **Supplementary Figures 1-3; Supplementary Tables 2-4**).

### Control for genetic confounding using only GWAS summary statistics

When individual-level data for *X* and *Y* are unavailable, it is generally considered impossible to quantify the association between *X* and *Y*, let alone controlling for genetic confounding. Here, we introduce PENGUIN-S to estimate exposure-outcome association using only GWAS summary statistics. The key insight here is that when the two input GWAS summary statistics datasets (for *X* and *Y*) share a substantial subset of study participants, it is possible to estimate *ρ*_XY_, i.e., the phenotypic covariance between *X* and *Y*, from these GWAS summary data using the intercept of bivariate LD score regression^12^:

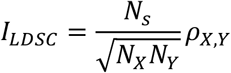

where *I*_*LDSC*_ is the intercept in bivariate LD score regression; *N*_X_, *N*_Y_, and *N*_S_ are the corresponding sample sizes of GWAS of *X*, GWAS of *Y*, and the number of shared samples between them. Therefore, we can replace 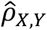 by 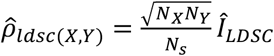 when estimating b:

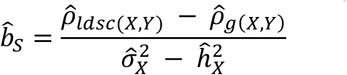

The other three terms (i.e., 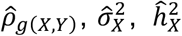) can be similarly obtained from GWAS summary statistics as demonstrated previously.

We can compute the standard error of PENGUIN-S estimate 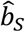 as follows:

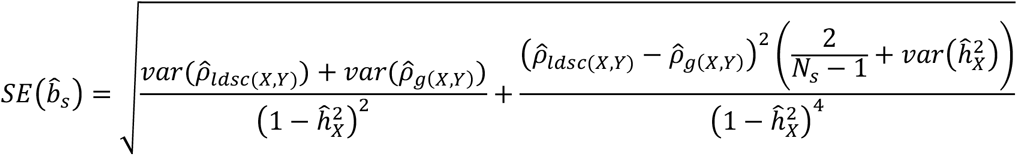

where 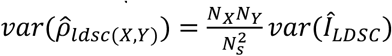 is obtained based on Jackknife estimation of 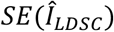 provided by bivariate LD score regression^12^.

### Simulation settings

We conducted simulations using real genotype data from WTCCC^43^. The WTCCC dataset contains 15,918 samples with 393,273 genotyped autosomal SNPs. We excluded individuals with genotype missing rate higher than 0.01 and removed SNPs matching any of the following conditions: (i) having minor allele frequency less than 0.05, (ii) having missing rate higher than 0.01 among all subjects, (iii) having Hardy-Weinberg Equilibrium p-value lower than 1e-6, and (iv) not found in 1000 Genomes Project Phase III European samples^78^. After quality control, 292,470 variants and 15,567 samples remained in the analysis.

We adapted the following data generating model to simulate exposure *X* and outcome *Y*. Let *G* denote a random vector of SNP genotype, and **α, β**, and **γ** are mutually independent SNP effects for genetic confounding, unconfounded genetics of *X*, and unconfounded genetics of *Y*, respectively:

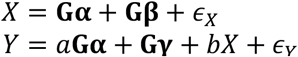

where *∈*_X_ and *∈*_Y_ are independent residuals, **𝒶**is a constant that controls the magnitude of genetic confounding, and *b* is the exposure effect we aim to estimate. True SNP effect sizes **α, β**, and **γ** follow point normal distribution with the proportion of causal variants denoted as **𝒫**. Based on this data generating model, we considered 3 simulation settings in total. In the first setting, there is no genetic confounding at all and *X* and *Y* are independent on both genetic and phenotypic levels. In the second scenario, the additive genetic component of *X*is the true genetic confounding factor. In the third and most general case, aggregated genetic confounding effects are part of *X*‘s genetics. Together, we generated distinct association structures between *X* and *Y* by varying distributions of SNP effects and magnitude of both genetic confounding effects and exposure effects:

***S****cenario* 1: **α** *is* **𝒫***oint mass at 0*, **β** *and* **γ** *are* **𝒫***oint normal, b* = *0*

***S****cenario* 2: **β** *is* **𝒫***oint mass at 0*, **α** *and* **γ** *are* **𝒫***oint normal, b* **∈**{*0, 0*.1, *0*.2, *0*.*3, 0*.4, *0*.5}

***S****cenario 3*: **α, β**, and **γ** are all point normal, *b* **∈** {*0, 0*.1, *0*.2, *0*.*3, 0*.4, *0*.5}

Detailed parameter specifications for simulation are summarized in **Supplementary Table 1**. We set the heritability of both *X* and *Y* to be 0.5 and simulated both sparse and polygenic genetic architecture by alternating **𝒫** between 0.1% and 5% in each setting. SNPs with non-zero effects were randomly selected across the genome and genetic components (i.e., PGS without measurement error) were calculated using PLINK^79^. To obtain trait values, we added simulated Gaussian noises such that the variance of phenotypes is 1 for both *X* and *Y*. For settings 2 and 3 where the exposure *X*has an effect on outcome *Y*, we obtained *X* first and calculated *Y* based on parameters specified in the true data generating model of *Y*. Each setting was repeated 100 times.

We compared PENGUIN and PENGUIN-S with five approaches: marginal regression between *X* and *Y* without controlling for genetic confounding, PGSX and PGSY which add the PGS of exposure *X* or outcome *Y* in the regression as a covariate, and GsensX and GsensY which apply structural equation model to remove measurement error in PGSX and PGSY^18^. For fair comparison, we partitioned the WTCCC datasets into 3 subsets for GWAS model training (training data; N=12,000), PGS calculation and outcome-exposure association testing (testing data; N=3,000), and calculation of LD scores (N=567). We performed GWAS on the training data to obtain summary statistics for both *X* and *Y* using PLINK^79^. For PGS and Gsens analyses, we used clumping and thresholding (C+T) PGS. We clumped GWAS summary statistics using an LD window size of 1Mb, *r*^**2**^ threshold of 0.1, and p-value cutoff of 1. For PENGUIN, we estimated the heritability of *X* and genetic covariance between *X* and *Y* using LD score regression^11,12^, and used the testing dataset to conduct statistical inference on b. PENGUIN-S only used GWAS summary statistics for *X* and *Y* obtained from the training data as input. We calculated and compared the coverage of 95% confidence intervals and proportion of p-value smaller than 0.05 across 100 replications from each method. Lastly, to investigate whether each method shows robust performance when there is a sample overlap between training and testing data, we considered two additional scenarios within each simulation setting. In the first scenario, 1500 individuals in the testing dataset were included in the GWAS training data (i.e., 50% overlap). In the second scenario, all 3000 testing samples came from the training data (i.e., 100% overlap). We repeated the analysis described above and compared each method’s performance under these two scenarios. All simulation results are summarized in **Supplementary Tables 2-4**.

### Estimating EA effect on complex traits in UKB

UKB genotype data was imputed to the Haplotype Reference Consortium reference^41^. We only included individuals of European ancestry in the analysis. We also excluded SNPs with minor allele frequency below 0.01, imputation *R*^**$**^ below 0.9, Hardy-Weinberg equilibrium test p-value below 1e-6, or missing genotype call rate greater than 2%. 408,193 samples and approximately 7 million variants remained after quality control. Following recent studies^48,49^, we curated a list of 26 complex traits to be included in association analysis. To estimate effects of EA on binary traits other than ‘ever smoking’, we randomly selected controls in the phenotype data by maintaining a 1:2 case-control ratio. For ‘ever smoking’^80^, we used all available samples in UKB since the number of cases is larger than controls. Detailed information for these phenotypes is summarized in **Supplementary Table 5**. All phenotypes were first residualized by regressing out effects of sex, year of birth (YOB), interaction between sex and year of birth, and top 20 genetic principal components (PCs) and then standardized in advance.

We compared PENGUIN with two approaches: marginal linear regression between EA and each outcome, and PGSX which adjusts for EA PGS in the regression. For PGSX, we used EA3 GWAS summary statistics excluding 23&Me and UKB (N=324,162)^47^ to compute C+T PGS for UKB participants. LD-clumping was implemented with the same set of parameters used in simulation study. PENGUIN used EA3 summary statistics excluding 23&Me (N=766,345) and recently published GWAS summary statistics for each outcome as inputs, except for 6 quantitative traits whose GWAS results were not publicly available. For those 6 traits, we performed GWAS in UKB using PLINK^79^ while adjusting for sex, YOB, sex-YOB interaction, and top 20 PCs. Information about each GWAS summary-level dataset is included in **Supplementary Table 5**. LD score regression was used to calculate EA heritability and genetic covariance between each phenotype and EA based on LD scores computed from 1000 Genome Project Phase III EUR samples^11,12,78^. For each binary trait, genetic covariance was calculated on the observed scale that matches its case-control ratio in individual-level UKB phenotype data^81^. We further applied PENGUIN-S to calculate EA effects on 15 quantitative traits using UKB GWAS summary statistics obtained from PLINK^79^, while adjusting for the same set of covariates. For fair comparison between PENGUIN-S and PENGUIN, we did not perform any genomic control to GWAS summary statistics since doing so would change the scale of heritability and genetic covariance estimation. Additionally, we analyzed EA’s effect on SCZ and BD, respectively, while controlling for the cognitive and non-cognitive genetic components of EA separately. Publicly available cognitive and non-cognitive EA GWAS summary statistics were obtained from a previous study^54^.

### Multi-generational confounding control and estimating parental effects on offspring EA

We extended the PENGUIN framework to control for genetic confounding between two generations. Using maternal effect estimation for illustration, let **G**_*>M*_ and **G**_*o*_ denote maternal and offspring genotypes, and *M* and *0* denote standardized maternal and offspring phenotypes. Then we have:

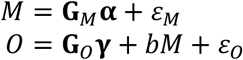

where *ε*_*M*_, and *ε*_*O*_ are independent residuals with mean 0, **α** and **γ** are SNP effects, and *b* is the maternal effect parameter of interest. *b* can be denoted as follows when the sample size approaches infinity:

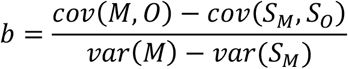

Here, ***S***_*M*_ is the additive genetic components underlying *M*. But an important distinction with the single-generation PENGUIN framework is that ***S***_*o*_ is the additive genetic component for how maternal genetics affect offspring outcome (i.e., the infinite-sample fitted value when regressing *0* on **G**_*M*_). In other words, ***S***_*o*_ is the infinite-sample PGS based on a GWAS in which we regress offspring outcome on maternal SNP data^82^. Then, we can replace each term with their respective estimator:

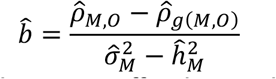

where 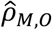 is the sample covariance between offspring and maternal phenotypes, 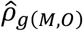 is the genetic covariance between offspring and maternal phenotypes, and 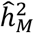 and 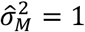 are the heritability and phenotypic variance of maternal phenotype. 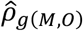 can be estimated from a standard GWAS of own phenotype and another GWAS of offspring phenotype (also referred to as GWAS-O and GWAS-M in our previous work^82^). Standard error and Z-statistics calculations remain the same for this two-generation model.

HRS genotype data was imputed based on 1000 Genomes Project reference data and accessed through NIAGADS. After removing SNPs with imputation quality score below 0.8, 15,567 samples and 18,144,468 SNP remained in the dataset. We obtained each HRS participant’s own, mother’s, and father’s years of schooling from the childhood family data. 12,068 HRS samples remained after matching participants with non-missing EA phenotypes and self-reported European ancestry. For each EA phenotype, we regressed out covariates including sex, YOB, and top 20 PCs and standardized the residuals prior to association analysis.

We compared PENGUIN with marginal regression and the PGS approach. Since only offspring genetic data are available, we could only apply PGSY which adjusts for offspring EA PGS in regression. We calculated C+T PGS using EA3 GWAS summary statistics excluding 23&Me, HRS, Add Health, and Wisconsin Longitudinal Study (N=742,903)^47^ on HRS samples using an LD window size of 1Mb, *r*^**2**^ threshold of 0.1, and p-value cutoff of 1. For PENGUIN, we applied LD score regression^11^ to estimate 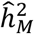 and 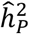 based on the same EA3 GWAS summary-level data used for PGS construction. To improve the GWAS sample size for parental genetic effect on offspring outcome, we used summary statistics from a previous GWAS in which we regressed offspring EA on either parent’s genotype (GWAS-MP; N=15,377; meta-analysis of UKB, WLS, and HRS)^82^. We estimated genetic covariance 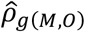 and 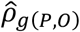 using this GWAS and the EA3 GWAS association statistics (N=742,903)^47^. As a secondary analysis where we relaxed the assumption that paternal and maternal SNPs have the same genetic effects on offspring outcomes, we replaced GWAS-MP by GWAS-M (regressing offspring EA on maternal genotype; N=3,477) and GWAS-P (regressing offspring EA on paternal genotype; N=1,560) based on UKB mother-offspring and father-offspring duos when evaluating maternal and paternal EA effects. For both GWAS-M and GWAS-P, we performed GWAS using PLINK^79^ while adjusting for offspring sex, offspring year of birth, top 20 PCs, and parental genotype chip. Association results from PENGUIN, PGSY, and marginal regression are summarized in **Supplementary Table 10**.

## Code availability

PENGUIN software is freely available at https://github.com/qlu-lab/PENGUIN.

## Author Contributions

Q.L. conceived the study.

Z.Z. and Q.L. developed the statistical framework.

Z.Z. performed the statistical analysis.

X.Y. assisted in UKB and HRS data processing and analysis.

J.M. helped in generalization of statistical methodology.

Z.Z. and S.D. implemented the software.

S.B. assisted in result interpretation.

J.F. advised on social genomics issues.

Q.L. advised on statistical and genetic issues.

Z.Z. and Q.L. wrote the manuscript.

All authors contributed to manuscript editing and approved the manuscript.

## Acknowledgments

The authors gratefully acknowledge research support from National Institutes of Health (NIH) grant R21 AG085162, and support from the University of Wisconsin-Madison Office of the Chancellor and the Vice Chancellor for Research and Graduate Education with funding from the Wisconsin Alumni Research Foundation (WARF). This study makes use of summary statistics from many GWAS consortia. We thank many GWAS investigators for providing publicly accessible GWAS summary statistics. This research uses data from HRS (NIAGADS accession number NG00119.v1). The HRS is supported by National Institute on Aging Grants U01AG009740, RC2AG036495, and RC4AG039029 and is conducted by the University of Michigan. No direct support was received from grant U01AG009740, RC2AG036495, and RC4AG039029 for this analysis. This study makes use of data generated by the Wellcome Trust Case Control Consortium. A full list of the investigators who contributed to the generation of the data is available from www.wtccc.org.uk. Funding for the project was provided by the Wellcome Trust under award 076113, 085475 and 090355. We thank members of the Social Genomics Working Group at University of Wisconsin for helpful comments. This research has been conducted using the UK Biobank Resource under Application 42148.

