## Supplementary Figures for "Controlling for polygenic genetic confounding in epidemiologic association studies"

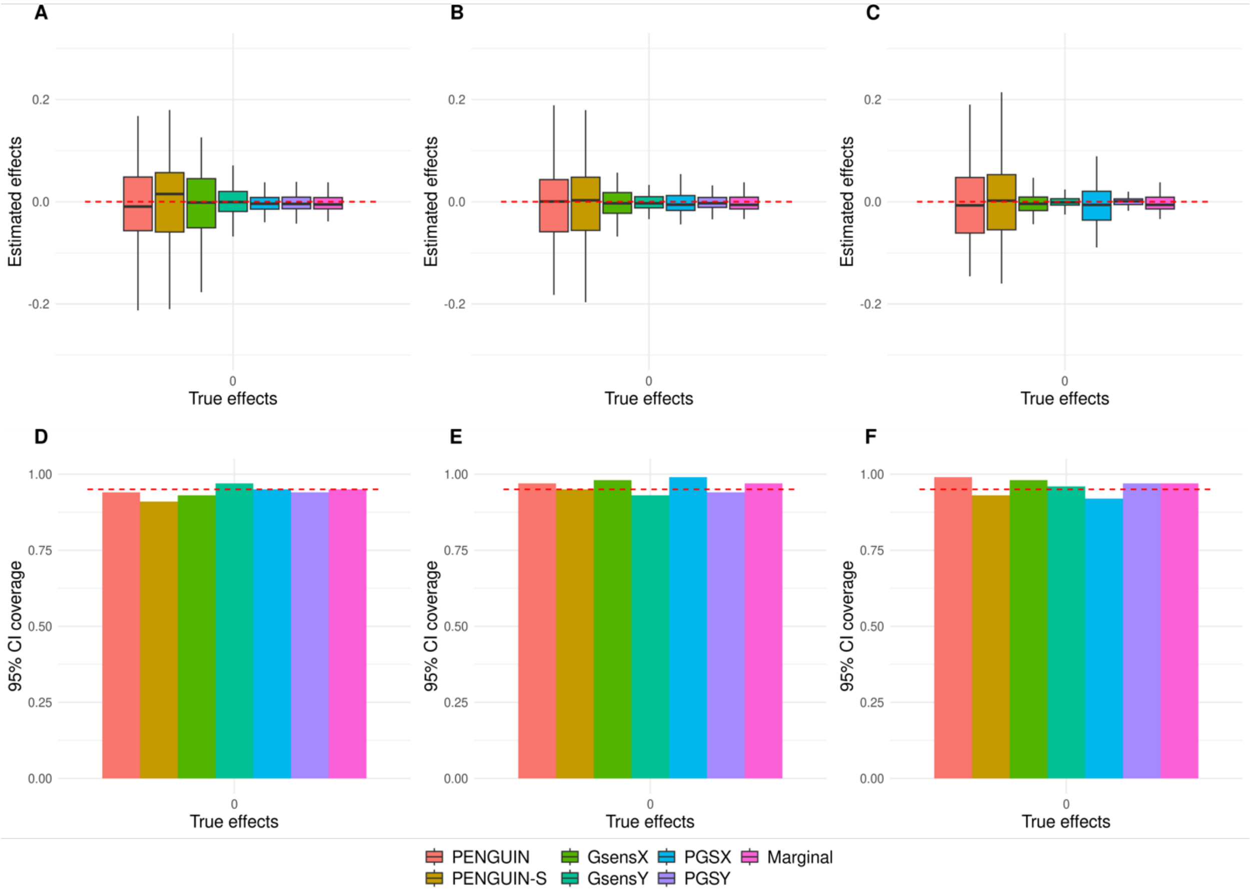


**Supplementary Figure 1. WTCCC simulation results in the sparse setting 1.** (**A**-**C**) Exposure-outcome associations estimated by different methods across 100 replications. (**D**-**F**) 95% confidence interval coverage for each method across 100 replications. GWAS summary-level data and individual-level testing dataset are independent in **A** and **D** (0% sample overlap), while 50% and all testing samples are included in the GWAS data in **B** and **E** (50% sample overlap) and **C** and **F** (100% sample overlap). Y-axis: exposure effects for **A-C** and 95% coverage for **D-F**; X-axis: true exposure effect size. Red dashed lines are true effects in **A**-**C** and 95% coverage thresholds in **D**-**F**. Across settings shown in this figure, the proportion of causal variants is 0.1% and there is neither genetic confounding nor exposure effect.

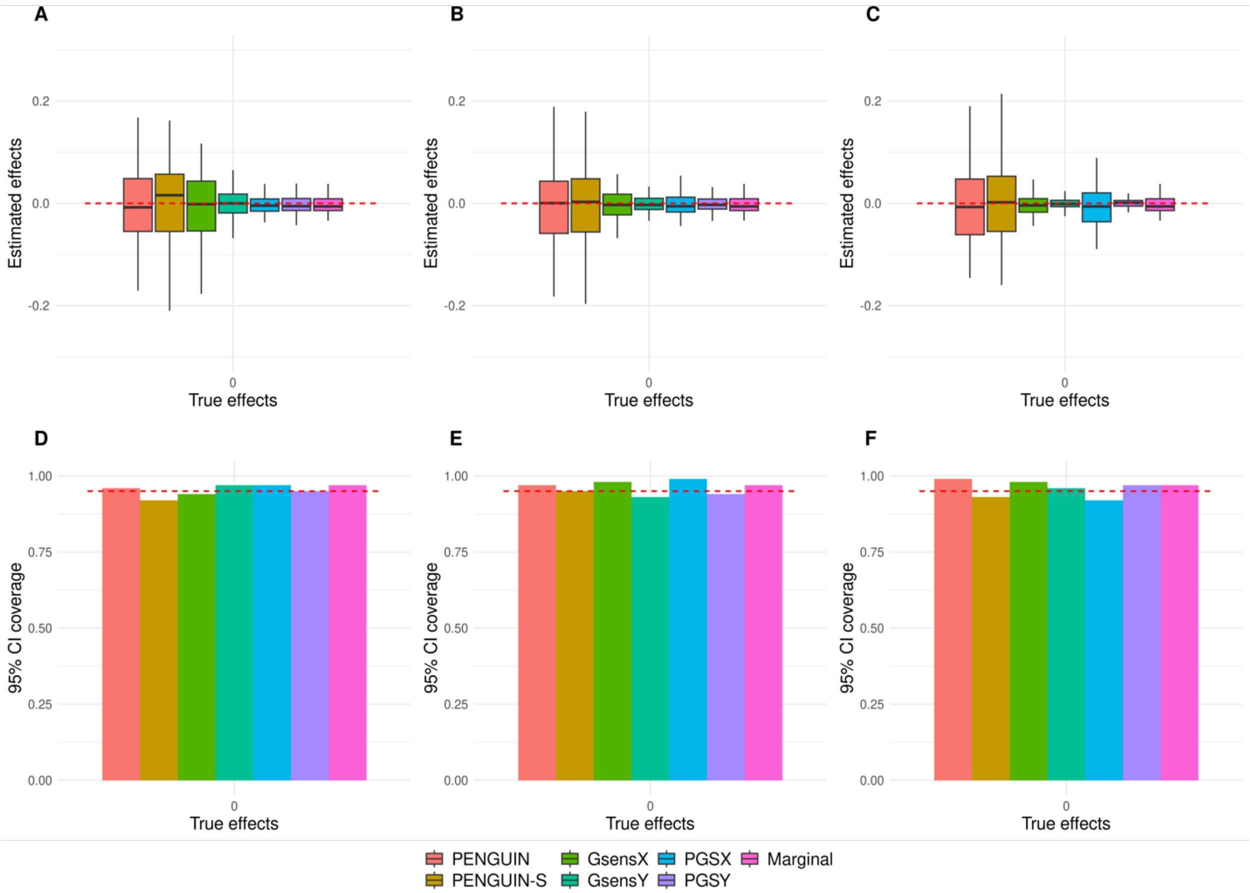


**Supplementary Figure 2. WTCCC simulation results in the polygenic setting 1.** (**A**-**C**) Exposure-outcome associations estimated by different methods across 100 replications. (**D**-**F**) 95% confidence interval coverage for each method across 100 replications. GWAS summary-level data and individual-level testing dataset are independent in **A** and **D** (0% sample overlap), while 50% and all testing samples are included in the GWAS data in **B** and **E** (50% sample overlap) and **C** and **F** (100% sample overlap). Y-axis: exposure effects for **A-C** and 95% coverage for **D-F**; X-axis: true exposure effect size. Red dashed lines are true effects in **A-C** and 95% coverage threshold in **D-F**. Across settings shown in this figure, the proportion of causal variants is 5% and there is neither genetic confounding nor exposure effect.

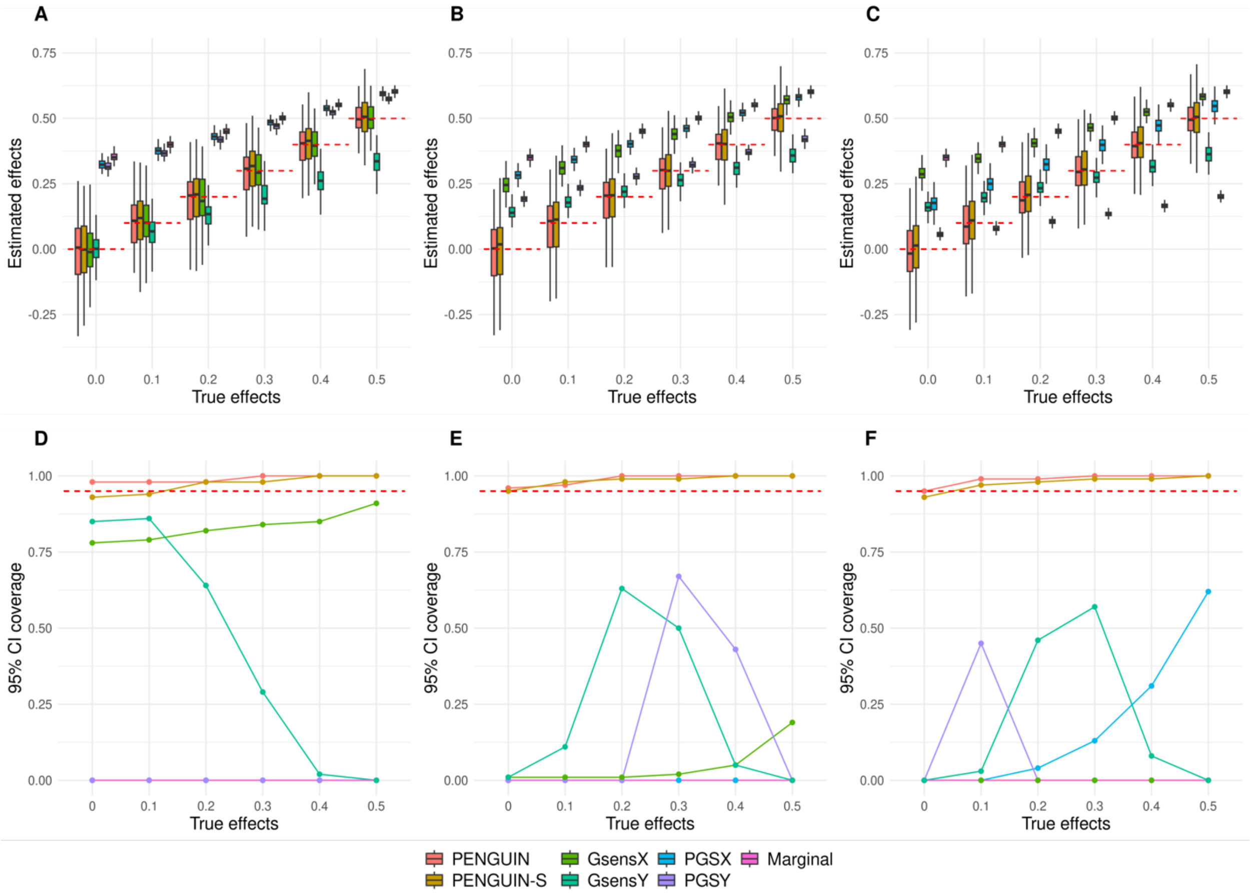


**Supplementary Figure 3. WTCCC simulation results in the sparse setting 2.** (**A**-**C**) Exposure-outcome associations estimated by different methods across 100 replications. (**D**-**F**) 95% confidence interval coverage for each method across 100 replications. GWAS summary-level data and individual-level testing dataset are independent in **A** and **D** (0% sample overlap), while 50% and all testing samples are included in the GWAS data in **B** and **E** (50% sample overlap) and **C** and **F** (100% sample overlap). Y-axis: exposure effects for **A-C** and 95% coverage for **D-F**; X-axis: true exposure effect size. Red dashed lines are true effects in **A**-**C** and 95% coverage threshold in **D**-**F**. Across settings shown in this figure, proportion of causal variants is 0.1% and true genetic confounding effects are the entire genetic component for the exposure.

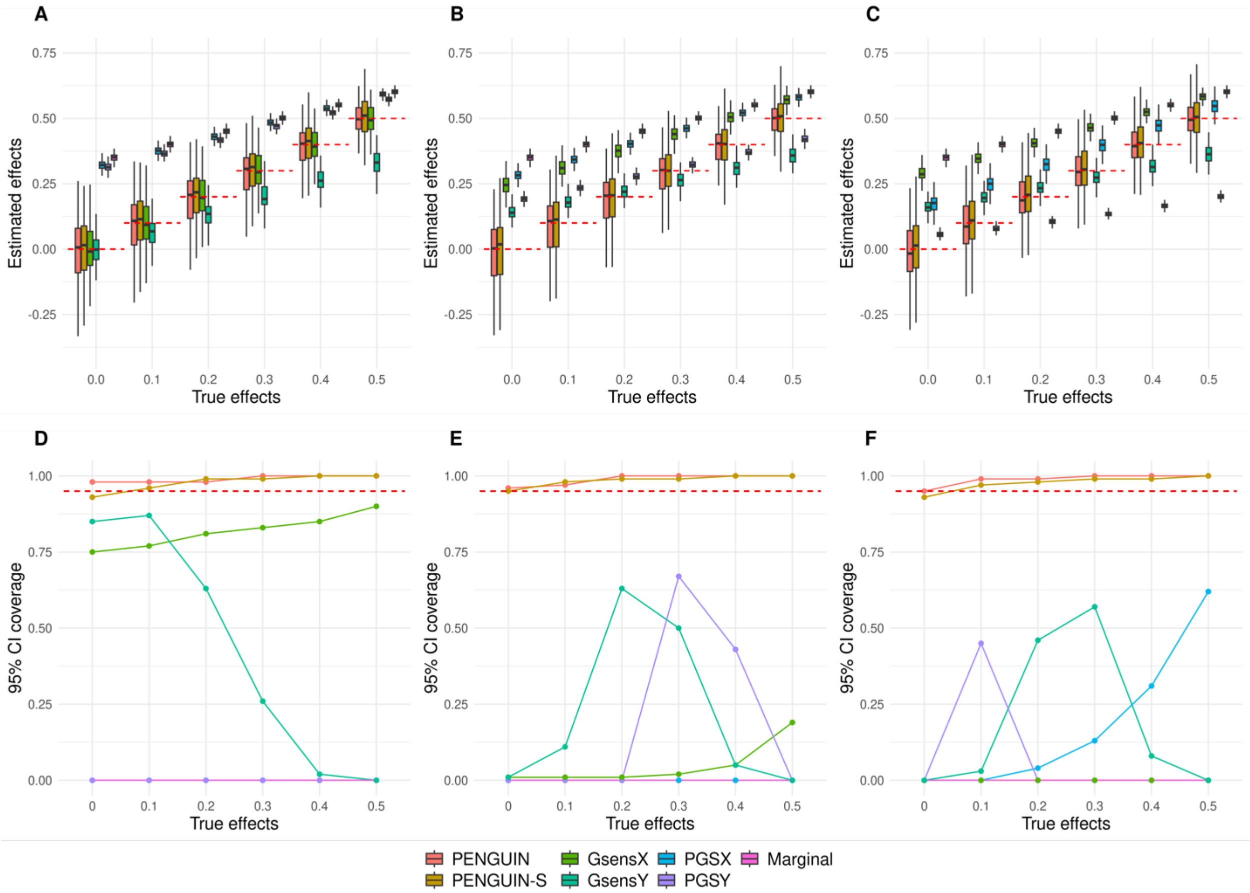


**Supplementary Figure 4. WTCCC simulation results in the polygenic setting 2.** (**A**-**C**) Exposure-outcome associations estimated by different methods across 100 replications. (**D**-**F**) 95% confidence interval coverage for each method across 100 replications. GWAS summary-level data and individual-level testing dataset are independent in **A** and **D** (0% sample overlap), while 50% and all testing samples are included in the GWAS data in **B** and **E** (50% sample overlap) and **C** and **F** (100% sample overlap). Y-axis: exposure effects for **A-C** and 95% coverage for **D-F**; X-axis: true exposure effect size. Red dashed lines are true effects in **A-C** and 95% coverage threshold in **D-F**. Across settings shown in this figure, proportion of causal variants is 5% and true genetic confounding effects are the entire genetic component for the exposure.

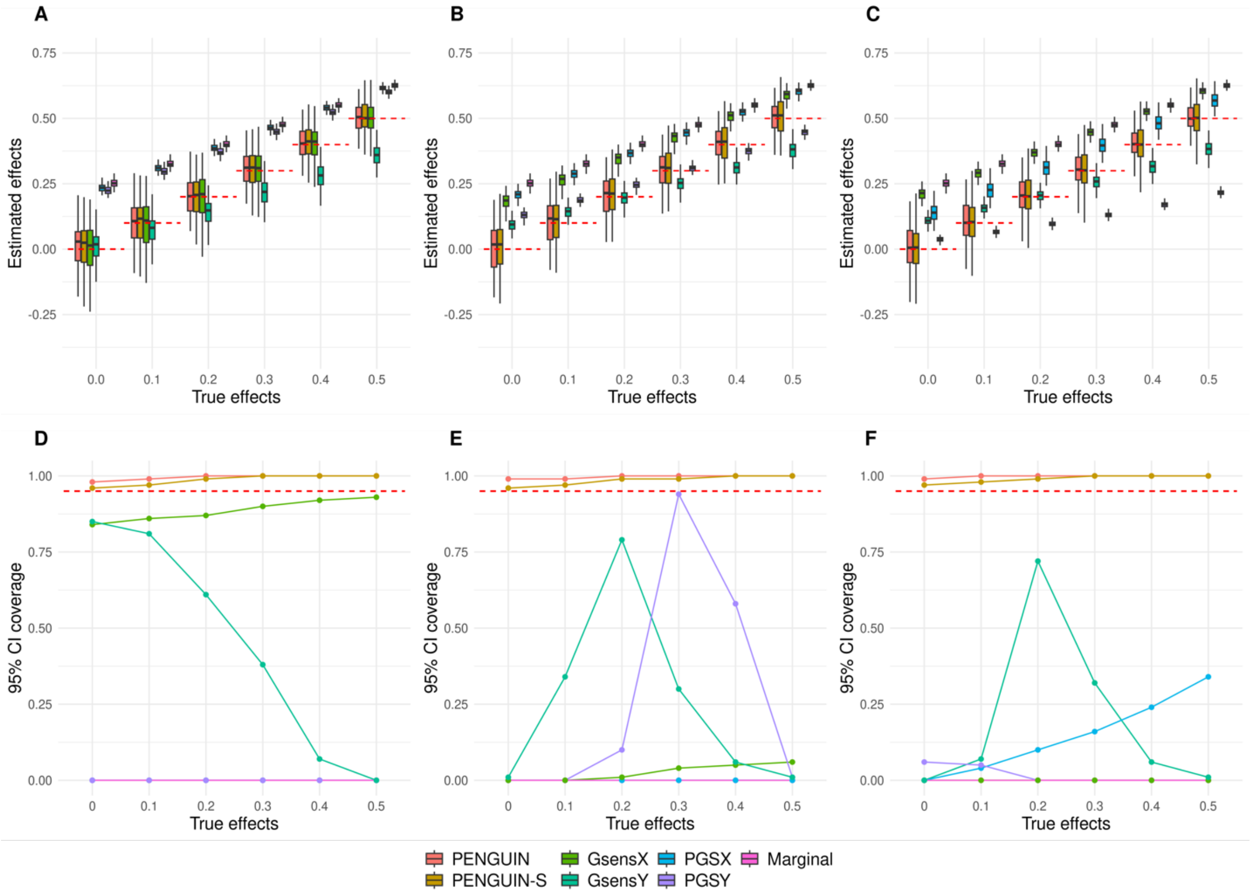


**Supplementary Figure 5. WTCCC simulation results in the sparse setting 3.** (**A**-**C**) Exposure-outcome associations estimated by different methods across 100 replications. (**D**-**F**) 95% confidence interval coverage for each method across 100 replications. GWAS summary-level data and individual-level testing dataset are independent in **A** and **D** (0% sample overlap), while 50% and all testing samples are included in the GWAS data in **B** and **E** (50% sample overlap) and **C** and **F** (100% sample overlap). Y-axis: exposure effects for **A-C** and 95% coverage for **D-F**; X-axis: true exposure effect size. Red dashed lines are true effects in **A**-**C** and 95% coverage threshold in **D**-**F**. Across settings shown in this figure, proportion of causal variants is 0.1% and true genetic confounding effects are part of the genetic component for the exposure.

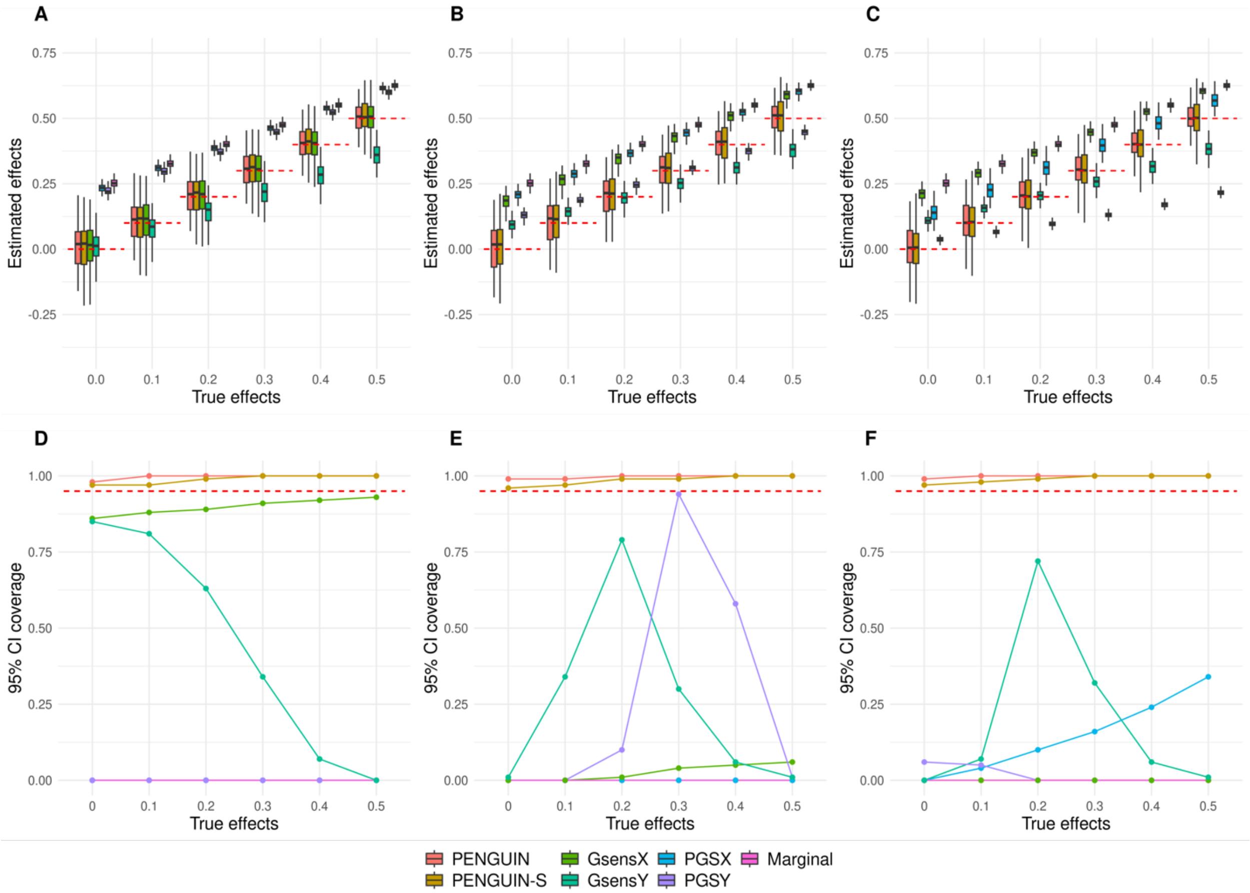


**Supplementary Figure 6. WTCCC simulation results in the polygenic setting 3.** (**A**-**C**) Exposure-outcome associations estimated by different methods across 100 replications. (**D**-**F**) 95% confidence interval coverage for each method across 100 replications. GWAS summary-level data and individual-level testing dataset are independent in **A** and **D** (0% sample overlap), while 50% and all testing samples are included in the GWAS data in **B** and **E** (50% sample overlap) and **C** and **F** (100% sample overlap). Y-axis: exposure effects for **A-C** and 95% coverage for **D-F**; X-axis: true exposure effect size. Red dashed lines are true effects in **A**-**C** and 95% coverage threshold in **D**-**F**. Across settings shown in this figure, proportion of causal variants is 5% and true genetic confounding effects are part of the genetic component for the exposure.

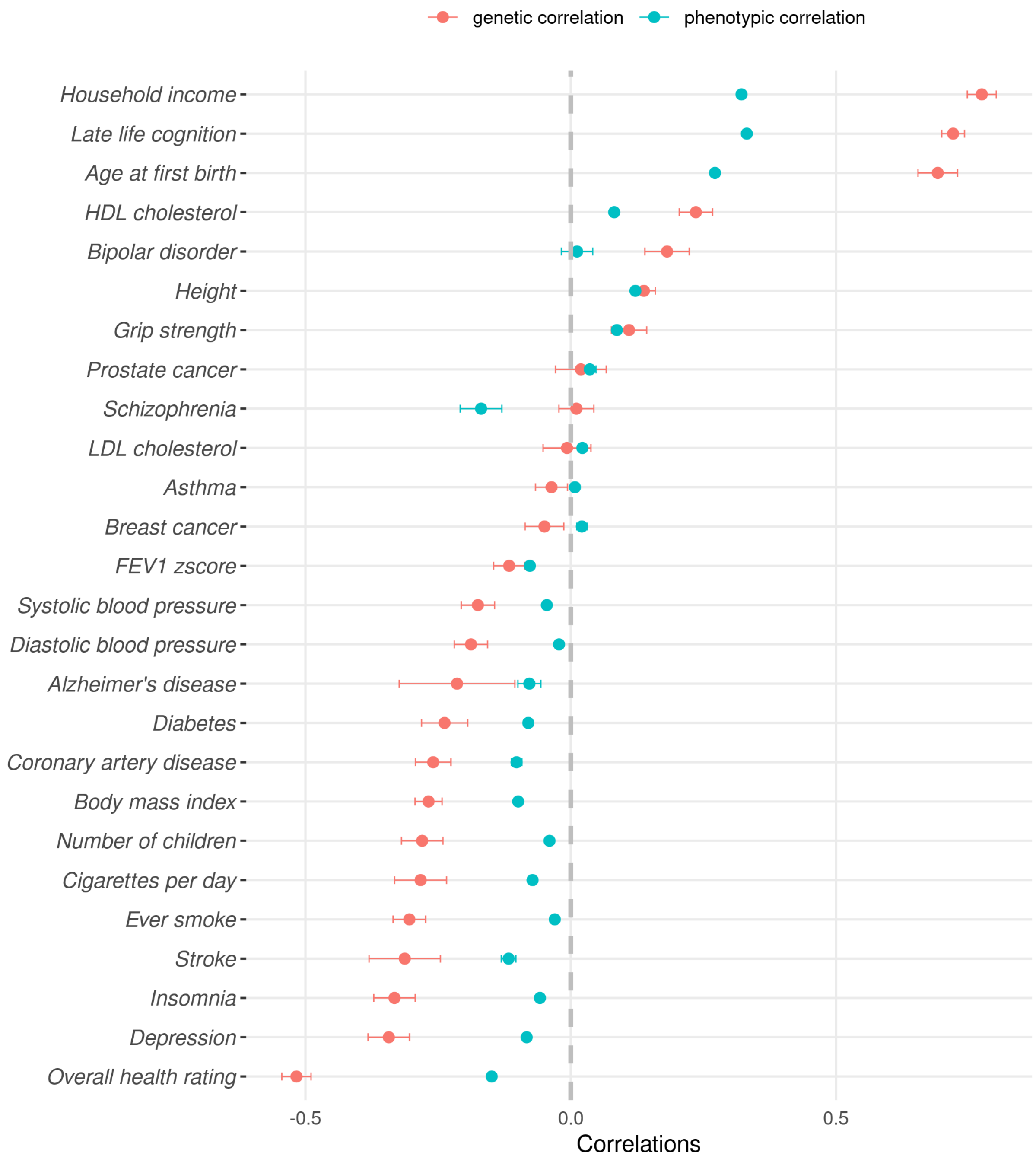


**Supplementary Figure 7. Phenotypic and genetic correlations between complex traits and EA in UKB**. Phenotypic correlations and standard errors are calculated using individual-level UKB phenotype data. Genetic correlations and standard errors are computed by LD score regression using GWAS summary statistics. Y-axis: outcome traits; X-axis: correlations.

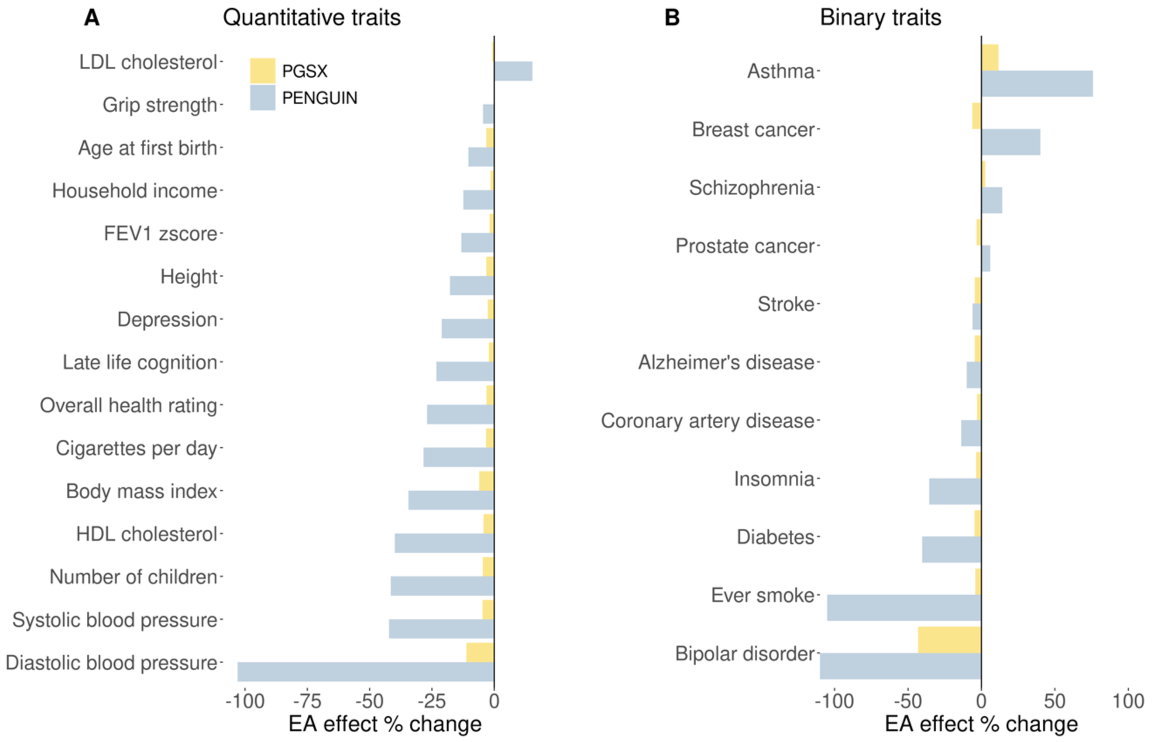


**Supplementary Figure 8. Relative changes in EA effects on various complex traits in UKB**. Compare EA effect estimates from PENGUIN and PGSX, using marginal regression coefficients as the baseline. Percentage changes in EA effect size point estimates for quantitative and binary traits are shown in panels **A** and **B**, respectively. Y-axis: outcome trait; X-axis: percentage change in EA effect size estimates (capped at -100% to the left end of X-axis).

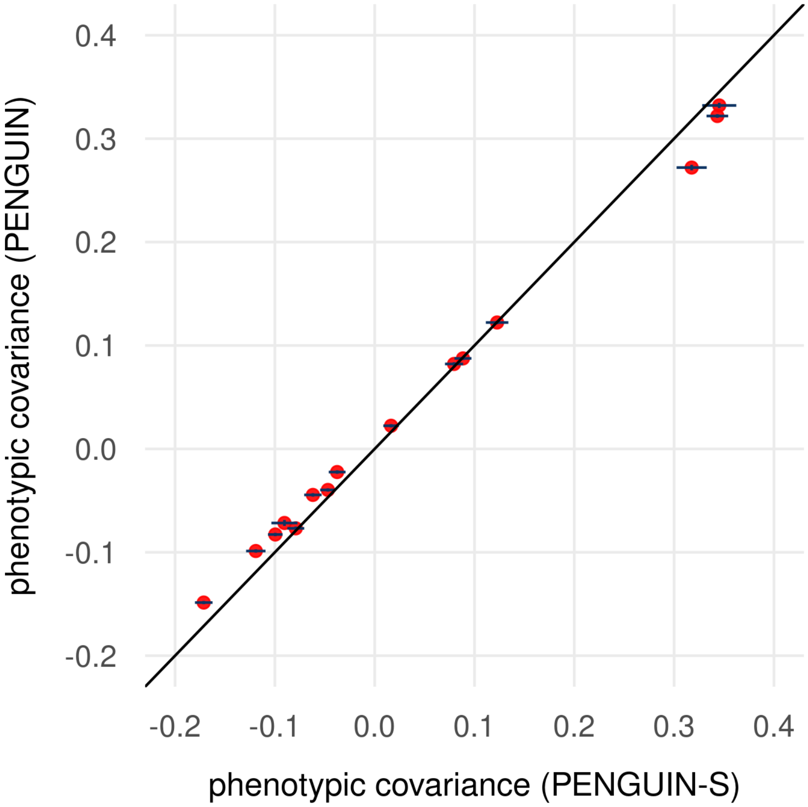


**Supplementary Figure 9. Comparison between phenotypic correlations** estimated **by PENGUIN and PENGUIN-S for 15 quantitative traits**. Y-axis: PENGUIN estimates; X-axis: PENGUIN-S estimates. The black solid line is the diagonal line. Standard error bars from both approaches are shown for all data points.

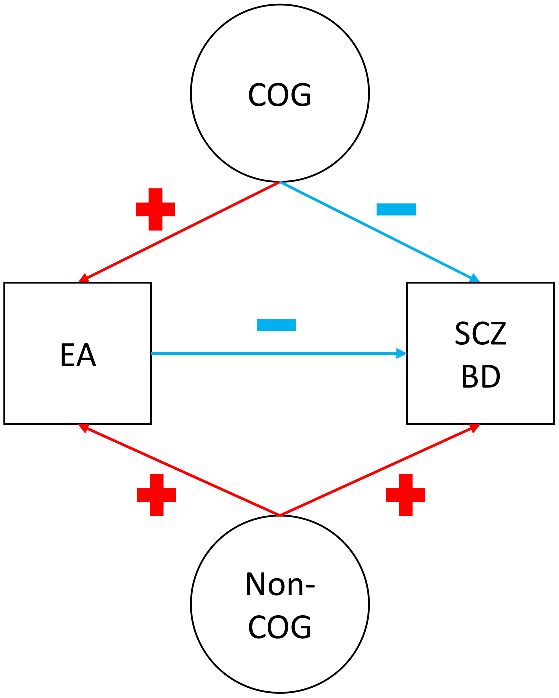


**Supplementary Figure 10. An illustration of genetic confounding between SCZ/BD and EA.** Phenotypes are shown in squares and genetic confounding effects are shown in circles. Red: positive effects; blue: negative effects. SCZ: schizophrenia; BD: bipolar disorder; EA: educational attainment.

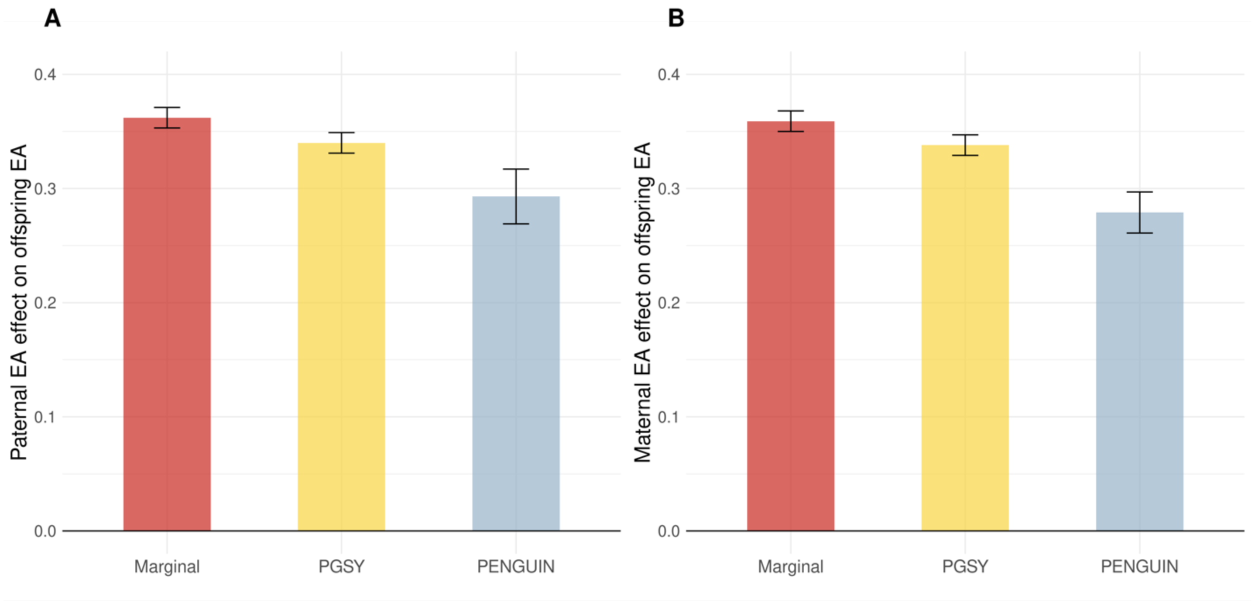


**Supplementary Figure 11. Effects of parental EA on offspring EA.** Paternal EA (**A**) and maternal EA (**B**) effect sizes are estimated from PENGUIN, PGSY, and marginal regression. EA GWAS-M and GWAS-P summary-level datasets based on UKB data (**Methods**) are used for estimating maternal and paternal EA effects, respectively. Y-axis: EA effect size estimates; X-axis: association approaches.
